# cloneMap: a R package to visualise clonal heterogeneity

**DOI:** 10.1101/2022.07.26.501523

**Authors:** Alexander M Frankell, Emma Colliver, Nicholas Mcgranahan, Charles Swanton

## Abstract

Genomic evolution and clonal heterogeneity in the soma are widespread and are involved in various pathologies with devastating consequences to human health, most prominently in neoplasia. Tools to interrogate and visualise the diversity of clonal populations in malignant or non-malignant tissues are currently lacking. cloneMap is a tool to represent clonal heterogeneity in a sample or set of samples, accounting for the clone size and phylogenetic relationships between clones. We apply cloneMap to several publicly available cancer sequencing datasets to demonstrate its ability to provide biological inferences of clonal heterogeneity. cloneMap is an R package available on github (https://github.com/amf71/cloneMap).

## Background

In the recent years it has become clear that genetic variation and somatic evolution is pervasive throughout the human body both in apparently healthy tissues^1–5^ and in various pathologies where it tends to be even more extensive^6–11^. While homogenising selective sweeps are common in pathologies such as neoplasia, continuous mutagenesis and selection drive subclonal diversification and heterogeneity both between samples from the same tissue and within single samples. This heterogeneity has important role a oncogenesis^10,11^, immune recognition^12,13^, aging^1,5^ and other forms of tissue damage^3,7^ and is now forms a large field of research worldwide.

Tools to perform subclonal deconvolution and phylogenetic tree building are now common^14–18^ however, while phylogenetic trees provide information about clonal relationships, they do not provide information about the relative clone sizes, *i*.*e*. cancer cell fractions (CCFs), and hence can obscure the extent or lack of clonal diversity in a given sample. Although some methods exist to present clone sizes^19^ these do not account for phylogenetic relationships had hence high heterogenous samples are not accurately represented. Standardised methods to visually display and compare heterogeneity of clones across a tissue in an automated and flexible fashion, accounting for phylogenetic relationships are currently lacking. Here we introduce cloneMap a standardised R package for publication-ready depictions of clonal heterogeneity, accounting for phylogenetic relationships between clones within a sample or set of samples. cloneMap is available on github (https://github.com/amf71/cloneMap).

## Methods & implementation

The cloneMap package uses a rasterised matrix to model relative clonal growth of sister clones at each level of the phylogenetic tree and produce semi-randomised, but reproducible, plots of the size and position each clone within a tissue. Sister clones grow together in intervals, respecting each other’s boundaries and those of the parent clone, to produce plausible arrangements of clonal structure. Nucleation of clones is semi-randomised, with nucleation away from other clones and the parent clone edge preferred. This provides realistic visual representation of diversity when visualising clonal structure of many samples at once. Required inputs are phylogenetic tree relationships and clone sizes (CCFs) which can be extracted from next generation sequencing data using various well established tools^14–16,18^. To enable reproducibility in the semi-randomised clone positions, an option to output the clone positions as a ‘cloneMap’ object is provided. This object can be saved and published to enable complete replication of an image. Plotting from a ‘cloneMap’ object also provides a significant processing time improvement for replotting the same image in several instances. CCFs can be provided as fractions or percentages and are adjusted to the CCF of the clonal cluster provided. CCFs are also adjusted to be phylogenetic tree-consistent, *i*.*e*. a clone cannot have a CCF less than the sum of its daughter clones, which can occur to due technical noise in the CCF measurements. This is performed from the top of the tree recursively, decreasing the CCF of daughter clones to fit within parents, and is accompanied by an informative message about CCF adjustments. Clone colours can be specified manually using a table of clones and colours or using a palette from the RColourBrewer package. Providing the same clone colour table to several cloneMaps from the same tissue allows the user to easily match clone colours across plots. CloneMap also allows unrooted trees, i.e. where some clones are unrelated, sharing no mutations, such as is common in data derived from normal tissues^7^. When this is the case CCFs become a relative measure of clone size and the overall percentage of mutated cells in the tissue can be specified and represented accurately. Border thickness, border colour, the extent of smoothing of clone edges and resolution of the clone positions can also be specified. The cloneMap package is dependent on a number of R packages: qlcMatrix, raster, sf, RColourBrewer, rgeos, smoothr and parrellel.

## Results

We applied the cloneMap method to two publicly available datasets, TRACERx a prospective cohort of 100 multi-region sequenced non-small cell lung cancers from Jamal-Hanjani et al^10^ and a cohort of seven multi-region sequenced ovarian high grade serous carcinomas from McPherson et al^20^.

Figure 1A & B demonstrates the additional information provided by the cloneMap depiction beyond a phylogenetics tree using two example tumours from the TRACERx lung cancer dataset. While both phylogenetic trees seem similarly diverse, we can see the diversity of clonal prevalence is highly distinct. Figure 1A depicts a highly diverse mixture of subclonal genotypes whereas Figure 1B shows a tumour dominated by the clonal genotype and contains only a small minority of cells conferring subclonal genotypes. The cloneMap package may be applied to each sample from a patient separately, as shown for an ovarian tumour in figure 1C, to represent the clonal heterogeneity between regions of the same tissue. We can see that some clones are highly regionally localised whereas others tend to spread across several regions which may indicate differences in growth rates or capacities for migration. Figure 1D shows a single cloneMap representing the same case which allows comparisons of diversity among a cohort of tumours (all ovarian tumours are shown in supplementary figure 1). The TRACERx 100 cohort is represented using cloneMaps in figure 1E, separated by stage. While there is a range of clonal diversities across all groups, there is a clear, visible increase in diversity in the later stage groups. We also observe differences in clonal diversity relative to genome doubling status (supplementary figure 2A) and histology (supplementary figure 2B).

**Figure 1.**
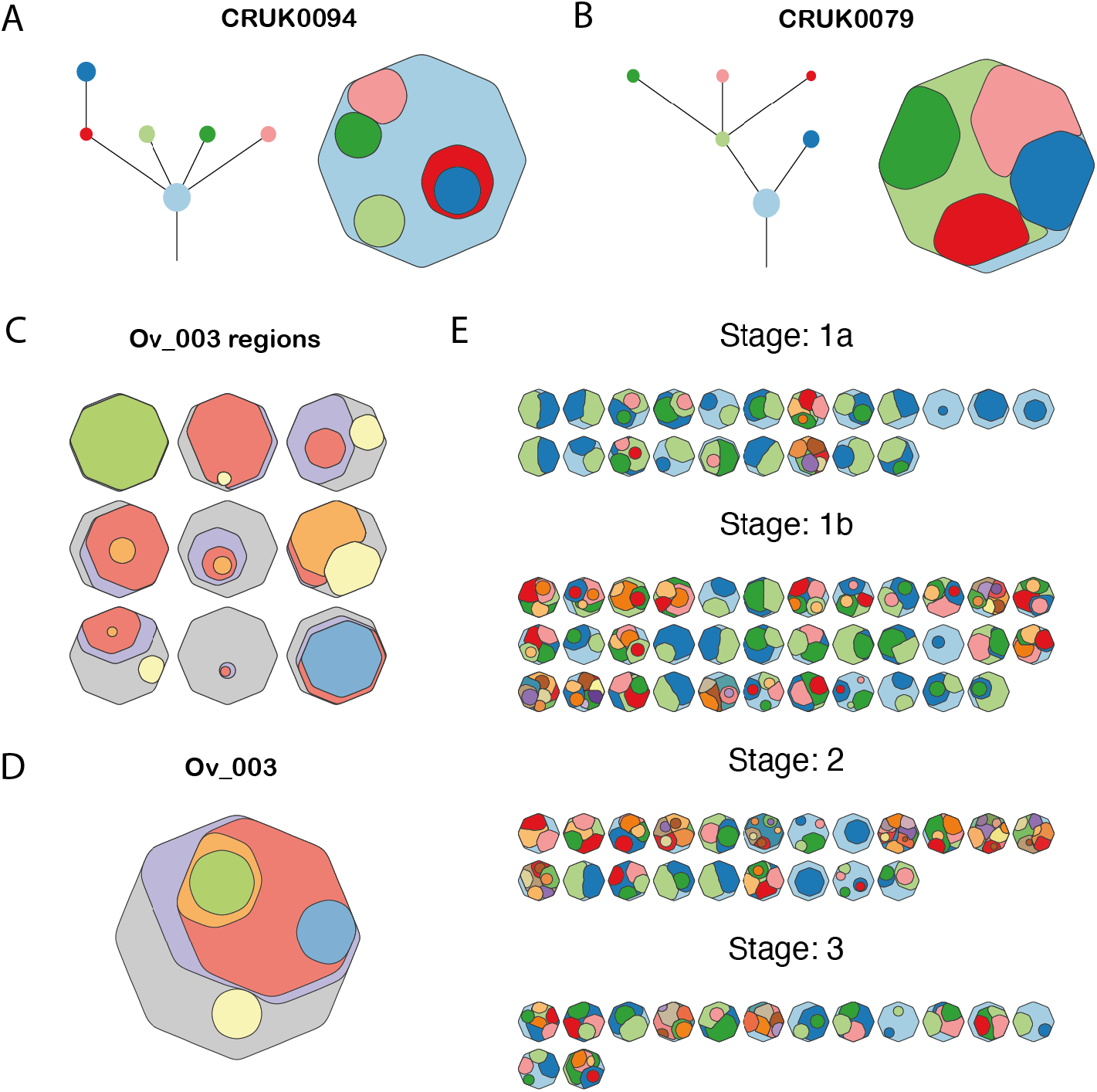
Utility of the cloneMap package to visualise tumour heterogeneity. **A**. A lung tumour with a moderately branched phylogenctic tree and low diversity of clonal prevelence. **B**. A lung tumour with moderately branched phylogenetic tree and high diversity of clonal prevelence **C**. An ovarian tumour were clonal prevelence shown per region. **D**. The same ovarian tumour with clonal prevelence summarised in one cloneMap. **E**. CloneMaps showing clonal prevelence for the TRACERx 100 Non-small-cell lung cancer cohort seperated by clinical stage.

The code to reproduce figures included in this article, from data from download to figure drawing, is provided within the associated R package (https://github.com/amf71/cloneMap).

## Conclusions

The studies of somatic evolution and heterogeneity are growing rapidly in number and ambition. These studies must be equipped with tools to analyse and interrogate the resulting data so that critical inferences are captured. CloneMaps is a tool to allow rapid and reproducible depictions of clonal frequencies, accounting for phylogenetic relationships, so as to allow comparisons of genetic structure and diversity across samples from the same tissue or tissues from difference individuals. Such insights will be vital in fields like oncology where clinical decision making may rest on such inferences.

---

**Supplementary figure 1.**
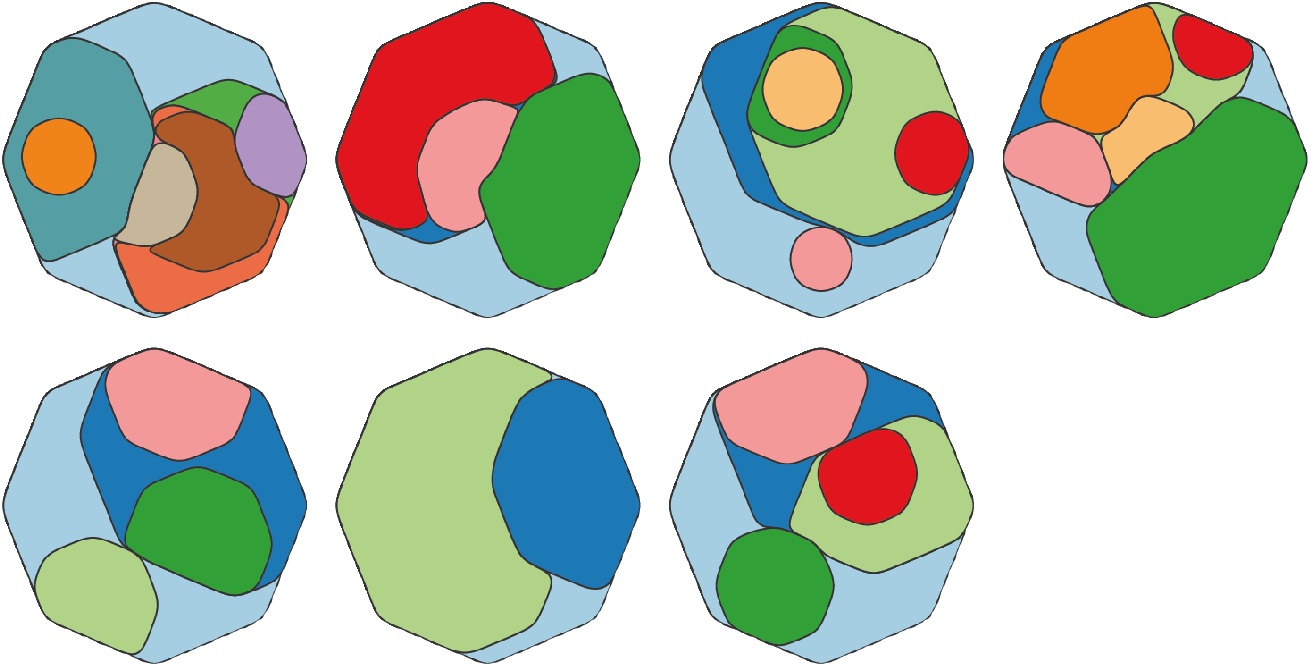
CloneMaps for 7 Ovarian tumour from McPherson et al showing a veriety of clonal compositions.

**Supplementary figure 2.**
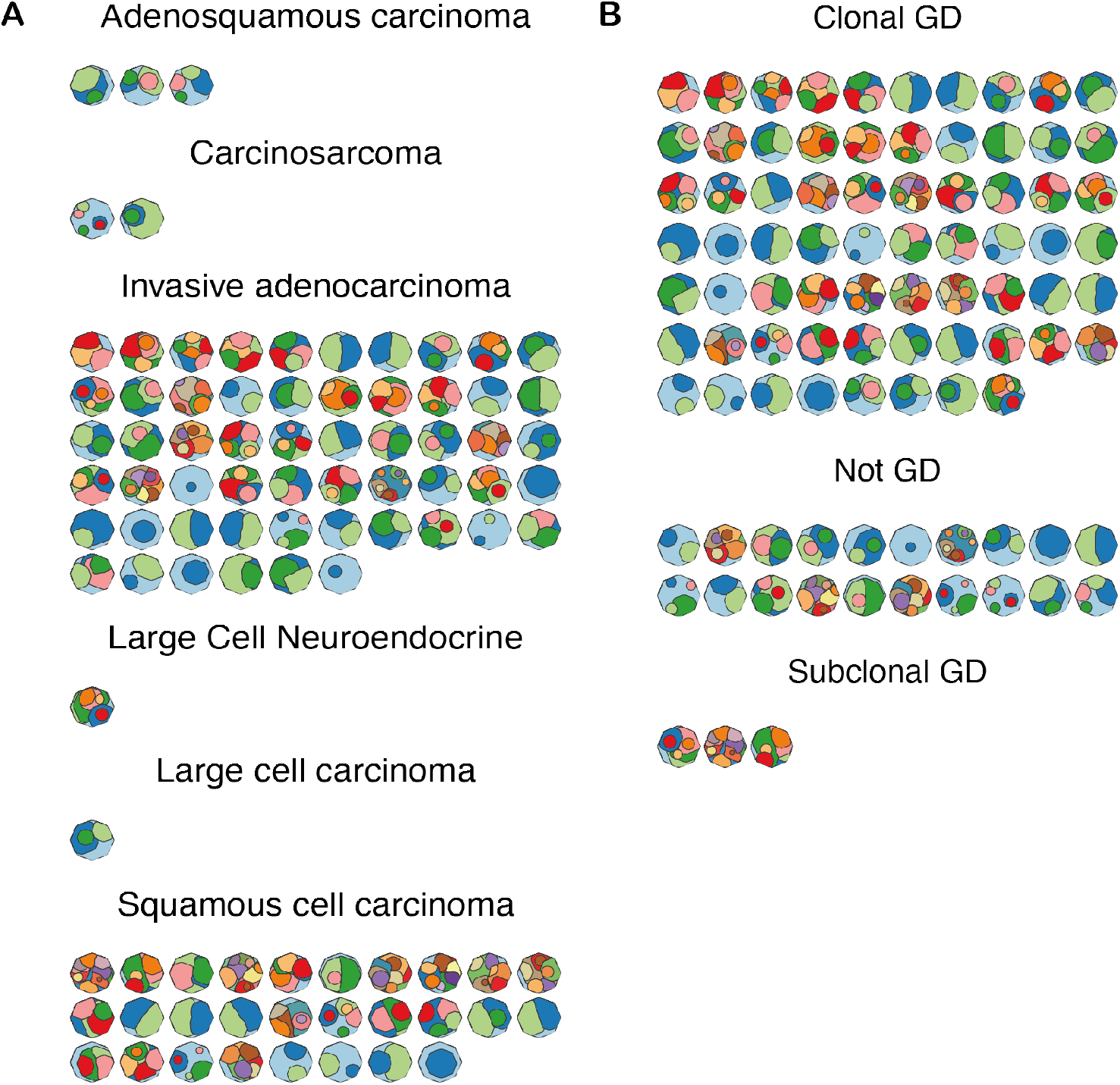
CloneMaps for the TRACERx 100 cohort. **A**. Tumours seperatered by histoogy. **B** Tumour seprated by genome doubling (GD) status

## Bibliography

1. Martincorena, I. et al. High burden and pervasive positive selection of somatic mutations in normal human skin. Science (80-.). 348, 880–886 (2015).

2. Lee-Six, H. et al. Population dynamics of normal human blood inferred from somatic mutations. Nature (2018) doi:10.1038/s41586-018-0497-0.

3. Yoshida, K. et al. Tobacco smoking and somatic mutations in human bronchial epithelium. Nature (2020) doi:10.1038/s41586-020-1961-1.

4. Lee-Six, H. et al. The landscape of somatic mutation in normal colorectal epithelial cells. Nat. 2019 5747779 574, 532–537 (2019).

5. Martincorena, I. et al. Somatic mutant clones colonize the human esophagus with age. Science (80-.). eaau3879 (2018) doi:10.1126/SCIENCE.AAU3879.

6. Olafsson, S. et al. Somatic Evolution in Non-neoplastic IBD-Affected Colon. Cell 182, 672-684.e11 (2020).

7. Brunner, S. F. et al. Somatic mutations and clonal dynamics in healthy and cirrhotic human liver. Nat. 2019 5747779 574, 538–542 (2019).

8. Cross, W. et al. The evolutionary landscape of colorectal tumorigenesis. Nat. Ecol. Evol. 2, 1661–1672 (2018).

9. Ross-Innes, C. S. et al. Whole-genome sequencing provides new insights into the clonal architecture of Barrett’s esophagus and esophageal adenocarcinoma. Nat. Genet. 47, 1038–1046 (2015).

10. Jamal-Hanjani, M. et al. Tracking the Evolution of Non–Small-Cell Lung Cancer. N. Engl. J. Med. 376, 2109–2121 (2017).

11. Turajlic, S. et al. Deterministic Evolutionary Trajectories Influence Primary Tumor Growth: TRACERx Renal. Cell 173, 595-610.e11 (2018).

12. rosenthal, rachel et al. Neoantigen-directed immune escape in lung cancer evolution Heterogeneity of immune infiltration. Nature 8, 9.

13. Wolf, Y. et al. UVB-Induced Tumor Heterogeneity Diminishes Immune Response in Melanoma. Cell 179, 219-235.e21 (2019).

14. Roth, A. et al. PyClone: Statistical inference of clonal population structure in cancer. Nat. Methods 11, 396–398 (2014).

15. Miller, C. A. et al. SciClone: Inferring Clonal Architecture and Tracking the Spatial and Temporal Patterns of Tumor Evolution. PLoS Comput. Biol. 10, 1003665 (2014).

16. Malikic, S., McPherson, A. W., Donmez, N. & Sahinalp, C. S. Clonality inference in multiple tumor samples using phylogeny. Bioinformatics 31, 1349–1356 (2015).

17. Dang, H. X. et al. ClonEvol: Clonal ordering and visualization in cancer sequencing. Ann. Oncol. 28, 3076–3082 (2017).

18. Bolli, N. et al. Heterogeneity of genomic evolution and mutational profiles in multiple myeloma. Nat. Commun. 5, 1–13 (2014).

19. Birkbak, N. J. & Mcgranahan, N. Cancer Cell Review Cancer Genome Evolutionary Trajectories in Metastasis. (2020) doi:10.1016/j.ccell.2019.12.004.

20. McPherson, A. et al. Divergent modes of clonal spread and intraperitoneal mixing in high-grade serous ovarian cancer. Nat. Genet. 48, 758–767 (2016).

